# Somatic mutation landscape reveals differential variability of cell-of-origin for primary liver cancer

**DOI:** 10.1101/511790

**Authors:** Kyungsik Ha, Masashi Fujita, Rosa Karlić, Sungmin Yang, Yujin Hoshida, Paz Polak, Hidewaki Nakagawa, Hong-Gee Kim, Hwajin Lee

**Affiliations:** Biomedical Knowledge Engineering Laboratory, Seoul National University, Seoul 08826, South Korea.; Laboratory for Genome Sequencing Analysis, RIKEN Center for Integrative Medical Sciences, Tokyo 108-8639, Japan.; Bioinformatics Group, Department of Molecular Biology, Division of Biology, Faculty of Science, University of Zagreb, Horvatovac 102a, 10000 Zagreb, Croatia.; Liver Tumor Translational Research Program, Simmons Comprehensive Cancer Center, University of Texas Southwestern Medical Center, Dallas, TX 75390, USA.; Department of Oncological Sciences, Icahn School of Medicine at Mount Sinai, 1425 Madison Ave., NY 10029, USA.

**Keywords:** Cell-of-origin, Primary liver cancer, Whole genome sequencing, Epigenomics, Cancer genomics

## Abstract

**Background:** Primary liver tissue cancers display consistent increase in global disease burden and mortality. Identification of cell-of-origins for primary liver cancers would be a necessity to expand options for designing relevant therapeutics and preventive medicine for these cancer types. Previous reports on cell-of-origin for primary liver cancers was mainly from animal studies, and integrative research utilizing human specimen data was poorly established.

**Methods:** We analyzed a whole-genome sequencing data set for a total of 363 tumor and progenitor tissues along with 423 normal tissue epigenomic marks to predict the cell-of-origin for primary liver cancer subtypes.

**Results:** Despite the mixed histological features, the predicted cell-of-origin for mixed hepatocellular carcinoma / intrahepatic cholangiocarcinoma were uniformly predicted as a hepatocytic origin. Individual sample-level prediction revealed differential level of cell-of-origin heterogeneity depending on the primary liver cancer types, with more heterogeneity observed in intrahepatic cholangiocarcinomas. Additional analyses on the whole genome sequencing data of hepatic progenitor cells suggest these progenitor cells might not a direct cell-of-origin for liver cancers.

**Conclusions:** These results provide novel insights on the heterogeneous nature and potential contributors of cell-of-origin predictions for primary liver cancers.

## Background

Primary liver cancers (PLCs) is one of the major cancer types with increasing global disease burden over the years, reaching incidence rates and mortality over 900,000 per year (1, 2). This high morbidity and mortality of PLCs is due to the complex nature of the disease and lacking effective diagnostics and treatment besides multi-kinase inhibitors, thus strongly emphasizing the importance of relevant researches on early diagnosis and extensive drug development. In line with this, several endeavored researches were performed on identifying suitable diagnostic markers and targeted therapy-based treatments for PLCs, including the whole genome and exome-level profiling (3). So far, recent comprehensive efforts on investigating the genomics of PLCs revealed novel insights about the major mutation signatures, sub-classifications, and recurrent somatic mutations in coding regions (*TERT, TP53, CTNNB1, KRAS, IDH1/2*, etc.) and noncoding regions (*NEAT1* and *MALAT1*), which some of them are driver mutations and may associate with the clinical outcomes (4, 5). More investigations are underway to fully unveil the mechanisms and processes behind the progression of PLCs.

One of the complex, unanswered questions associated with the progression of PLCs is the possible cell-of-origins (COOs) corresponding to the various subtypes. PLC not only represents classical hepatocellular carcinoma (HCC) subtype, comprising of ~90% of PLCs, but also includes mixed hepatocellular and cholangiocarcinoma (Mixed) and intrahepatic cholangiocarcinoma (ICC), which are the two cancer subtypes displaying biliary phenotype with a different extent. COOs for these subtypes might depend on the location of a tumor within the liver and the differential clinical status associated with each tumor, represented by individual-level variability of cancer progression. So far, *in-vitro* and *in-vivo* experiments strictly at animal models proposed possible COOs for different subtypes of PLCs, including hepatocytes for HCCs, Mixed and ICCs, cholangiocytes for Mixed and ICCs, and bipotential hepatic progenitor cells (HPCs) for HCCs and ICCs (6). None of these are yet confirmatory due to the potential biases accompanied by cell cultures and genetic manipulation-based lineage-tracing animal model systems and lack of human level studies, and both evidences which indicates either differentiated cells or HPCs as a predominant COO for PLCs are present. For example, COO for HCCs were either reported as solely hepatocytes (7) or hepatocytes plus differentiated benign lesions derived from HPCs (8). For the COO for ICCs, hepatocytes which undergo conversion into cholangiocytes (9) or the billiary epithelial cells (10) were pointed out as possible options depending on the usage of different transgenic models. In addition, recent reports also suggest the possibility of de-differentiation of hepatocytes (7) and cholangiocytes (11) after the liver injury as potential sources of progenitor cells and PLCs, which further enhances the complexity of cellular origin for the liver cancer progression. Efforts on extrapolating these COO-related complexities by utilizing actual human cancer tissue data itself were scarce with one article partly visiting at a preliminary level (12), but no studies were yet performed in a fully comprehensive, inter-cohort manner. Thus, uncovering the major COOs matching to each subtype of PLCs and examining the potential variance of COOs across the tumors from different individuals remain highly necessary for the better understanding of the cancer progression for PLCs along with the early-stage diagnosis and possibly the treatment selection.

Here, we performed a computational approach to dissect out the possible COOs matching to each cancer subtype within PLCs and to interrogate possible individual tumor-level heterogeneity in COOs. For this, we analyzed the whole genome sequencing data from 320 of PLCs (256 HCCs, 8 Mixed, and 56 ICCs), 12 of extrahepatic biliary tract cholangiocarcinoma (BTCAs) based on the assumption that these cancer type would display predominant cholangiocytic COO, and 31 of HPCs and colon adult stem cells for assessing the possibility as a common COO for PLCs, along with 423 of chromatin features at the epigenome-level (see “Methods”). Our study not only confirmed the role of chromatin marks associated with possible COOs in shaping the mutation landscape of PLCs, but also uncovering the differential contribution of each COO in different subtypes of PLCs.

## Methods

### Data

For most analyses in this study, we used somatic mutation data of whole-genome sequencing (WGS) from the NCC-Japan liver cancer (LINC-JP), RIKEN-Japan liver cancer (LIRI-JP), and Singapore biliary tract cancer (BTCA-SG) projects after acquiring permission of ICGC (http://icgc.org). LINC-JP and LIRI-JP data consisted of a total of 282 samples with the exception of some cases which displayed multifocal or hypermutations, and these data were subgrouped according to the histological types (256 HCCs, 8 Mixed, and 18 ICCs). Data from BTCA-SG were all extrahepatic cholangiocarcinoma samples consisted of 12 samples without any particular subgroups.

The raw files of these datasets were analyzed along the standard GATK pipeline (https://www.broadinstitute.org/gatk/) and somatic mutations were called with the MuTect algorithm (http://archive.broadinstitute.org/cancer/cga/mutect) (13).

In addition to the data sets listed above, WGS-derived somatic mutation profile from additional 31 stem/progenitor samples (10 hepatic progenitor cells and 21 colon adult stem cells) and 38 ICCs from previous studies (5, 14) were utilized for the analysis related to hepatic progenitor cells (Fig. 3, Additional file 1: Figure S7) or as an independent cohort for predicting the COO of ICCs (Additional file 1: Figure S3) and assessing viral-infection associated COO predictions for ICCs (Additional file 1: Figure S6a). Somatic variants of these samples were called from a different method that was designed in each study comparing to the datasets we analyzed.

A total of 423 epigenomic data for chromatin feature selections, correlation analyses and COO prediction analyses was obtained from ENCODE (15) and NIH Roadmap Epigenomics Mapping Consortium (16). NIH Roadmap epigenomics data can be accessed through the NCBI GSE18927 in Gene Expression Omnibus site (https://www.ncbi.nlm.nih.gov/geo/). In addition, chromatin data for liver tissues derived from hepatitis virus infected patients (donor HPC8 and HPC17) was obtained from IHEC (https://epigenomesportal.ca/ihec/download.html).

To estimate the regional mutation density and average signal of chromatin features, autosomes were divided into each 1-megabase region except sectors containing low quality unique mappable base pairs, centromeres, and telomeres.

We calculated the frequency of somatic mutations and ChIP-seq reads in each 1-megabase region to figure out the regional mutation density and histone modification profiles. The value of DNase I peaks and replication was also used to calculate DNase I hypersensitivity and Repli-seq profiles in each 1-megabase region. All these calculations were performed using BEDOPS (17).

### Principal coordinate analysis

PCOA was employed to represent similarity/dissimilarity of mutation frequency landscapes among the samples. Each sample was represented in a two-dimensional space consisting of principal coordinates 1 and 2 using a dissimilarity matrix, which reflected Pearson correlation coefficient among the samples.

### Feature selection based on random forest algorithm

Our feature selection analysis applied a modified version used in the previous study (18). Briefly, training set of each tree was organized and the mean squared error and the importance of each variable were evaluated using out-of-bag data. To determine the ranking of importance for each variable, the values of each variable were randomly permuted and examined to each tree. The initial importance value of variable m was estimated by subtracting the mean squared error between the untouched cases and the variable-m-permuted cases. Eventually, the ranking of each variable was determined by averaging importance values of variable m in the entire tree. We constructed a total of 1000 random forest trees to predict regional mutation density from a total of 423 chromatin features and employed greedy backward elimination to pick out the top 20 chromatin marks. This method sequentially removed the chromatin marker with the lowest rank at each step. These random forest models were repeated 1000 times each. Generally, in our feature selection analysis, the mutation density was calculated by combining the samples corresponding to each cancer type.

### Prediction of cell-of-origin by grouping of chromatin features

To predict cell-of-origin (COO) for individual samples, chromatin marks were subgrouped based on the aggregate sample-level feature selection results. As a first step, we selected significant chromatin cell types above the cutoff score from the feature selection results using aggregated samples corresponding to each cancer type (Fig. 1a). Subsequently, we added relevant cell types and grouped the chromatin marks according to each selected cell type to evaluate the effect of cell-type specific chromatin on explaining variability of mutational landscapes among samples. For predicting the COO for HCCs, we simply utilized the importance ranking among variables from 423 chromatin features due to the fact that liver chromatin features were the only major type in the aggregated feature selection results for HCCs.

**Fig. 1.**
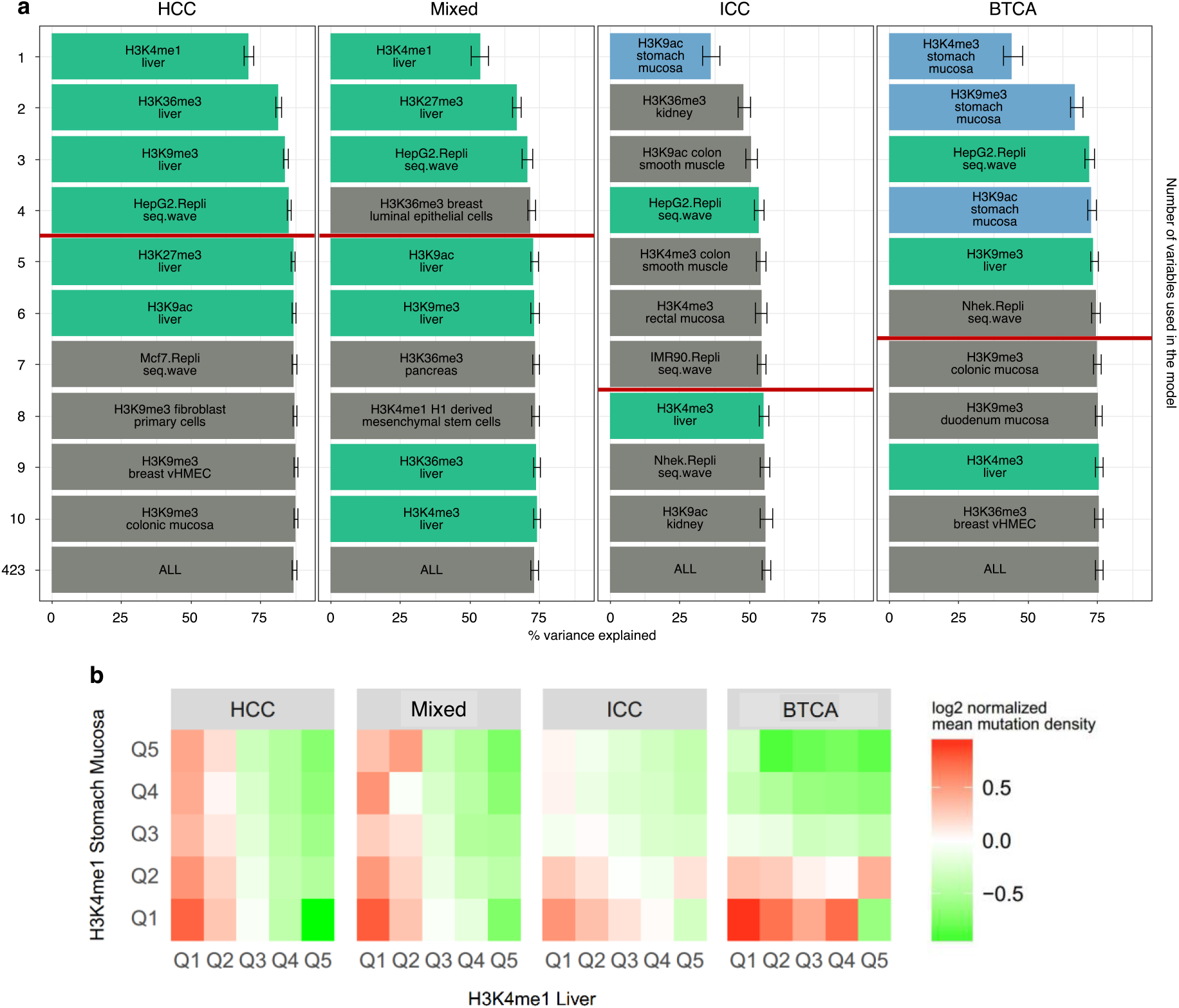
Cell-of-origin chromatin features delineating relations with the regional mutation frequency of HCCs, Mixed, ICCs and BTCAs. **a** Random forest regression-based chromatin feature selection using aggregated somatic mutation frequency data from HCC, Mixed, ICC and BTCA-SG samples. The rank of each chromatin feature is determined by importance values. The bar length represents the variance explained scores, and error bar shows minimum and maximum scores derived from 1,000 repeated simulations. Red lines represent the cutoff scores determined by the prediction accuracy of 423 features-1 s.e.m. Liver chromatin features are green-colored and stomach chromatin features are blue-colored. **b** Normalized mean mutation density per each PLC subtype and BTCAs plotted with respect to the density quintile groups of liver and stomach H3K4me1 marks.

### Signature analysis of mutational processes

Nonnegative matrix factorization (NMF) algorithm was employed to investigate mutation signatures as described in previous study (19). This methodology was utilized by factoring out frequency matrix of 96-trinucleotide mutation contexts from HCC, Mixed, ICC, BTCA-SG and HPC samples.

### Gene expression analysis

RNA-Seq experiments of HCC samples were performed previously (4), and the data had been deposited in the European Genome-phenome Archive. The reads were aligned onto the reference human genome GRCh37 using TopHat v2.1.1. Raw read counts per gene were computed using HTSeq with the GENCODE v19 annotation. Differential gene expression between hepatocytic- and non-hepatocytic-origin HCCs was analyzed using limma-voom v3.26.9 (20). Gene set enrichment analysis (GSEA) was performed using the GSEAPreranked v5 module on the GenePattern server (https://genepattern.broadinstitute.org).

## Results

### Aggregate Sample-level Correlations Between Chromatin Marks and Somatic Mutations of PLCs

Based on the previous findings about the close associations between the chromatin feature levels and regional variations in somatic mutation frequencies of tumor (18) and a number of precancerous lesions (21), we first hypothesized that the whole-genome mutation landscape of hepatocytic PLC subtype (HCCs) would exhibit closer relationship with the liver tissue (surrogate tissue for the hepatocytes) chromatin marks, whereas the mutation landscape of partial or fully biliary PLC subtypes (Mixed and ICCs) and the BTCAs would likely to display higher correlations with the chromatin marks from tissues containing either cuboidal or columnar epithelium (kidney, stomach or intestines as representative surrogate tissues for the cholangiocytes), depending on the extent of biliary phenotype and anatomical location. To examine differential associations among the mutation landscape for different subtypes of PLCs and the chromatin feature levels from normal tissues, we first employed a random-forest based feature selection method to identify the chromatin features responsible for explaining the possible variances in regional somatic mutation frequencies. To conduct the analysis, we utilized the 1-megabase window somatic mutation frequency data for three subtypes of PLCs (HCCs, Mixed and ICCs) and BTCAs at an aggregated sample level along with the 1-megabase window chromatin feature counts. As hypothesized, liver tissue chromatin marks served as major features with significance for HCCs, and stomach tissue chromatin mark served as the first-rank feature for ICCs and BTCAs (P < 2.2e-16, Mann-Whitney U-test between the first- and second-rank features of each PLC subtype; Fig. 1a). Surprisingly, liver tissue chromatin marks were major features explaining the regional mutation variation of Mixed albeit containing the biliary phenotype, implicating the unexpected skewness of possible COO towards to the hepatocytes for the particular subtype. The lower variance explained score for Mixed and ICCs comparing to the HCCs were at least in part likely due to the lower number of the samples and the total mutation load (Additional file 1: Figure S1a, b), indicating that the correlation between the liver tissue chromatin feature levels and the somatic mutation landscape of Mixed is similar to that of HCCs. In line with these result, spearman correlations between the regional mutation frequency of HCCs or Mixed and liver H3K4me1 chromatin mark level was the highest among different possible surrogate tissues, whereas stomach H3K4me1 chromatin mark level showed the highest correlation with the regional mutation frequency of BTCAs. (Additional file 1: Figure S2a, b). Spearman correlation values among the regional mutation frequency of ICCs and H3K4me1 of different tissues were overall low without any particular tissue type dependent differences, possibly due to both the lower mutation load and the possible variability in COOs which have been previously reported (12). Similar to the spearman correlation results, regional quintile-based mean mutation density data of HCCs and Mixed were relatively highly associated with the liver tissue H3K4me1 level comparing to the H3K4me1 level of stomach tissues, while mean mutation data of ICCs and and BTCAs display higher association towards the stomach tissue H3K4me1, with ICCs as a lesser extent (Fig. 1b). Collectively, these results demonstrate that the COO-associated chromatin features could delineate the relationships with the mutation landscape of PLCs and BTCAs.

### Individual Sample-level Cell-of-origin Predictions

To further assess the differential mutation landscapes and possible COOs of PLCs and BTCAs at the individual sample level, we conducted random forest algorithm-based COO analysis for each sample. This individual sample-based COO analysis exhibited the dominance of hepatocytic predicted COO for HCCs and Mixed, in contrast to the BTCAs which showed stomach tissues (a proxy tissue for extrahepatic cholangiocytes) possibly as a major COO (Fig. 2a). For ICCs, however, more heterogeneity of COO prediction was observed, and both hepatocytes and proxy tissues for cholangiocytes (kidney and stomach) were shown to be possible major COOs. This COO prediction pattern displayed consistency between different ICC cohorts (Additional file 1: Figure S3), thus emphasizing the heterogeneous nature of COO for ICCs. Our results not only replicated earlier findings on the COO of HCCs, ICCs and extrahepatic distal cholangiocarcinoma (DCCs) (12), but also additionally providing novel aspects about the complete predominance of hepatocytic predicted COO for Mixed tumors (8/8) and the implication of cuboidal cholangiocytes near the canal of hering (kidney tissue chromatin mark as a surrogate) could be another major COOs for ICCs besides the hepatocytes. In addition, 6 HCC samples showed non-hepatocytic predicted COO, thus inferring the possible distinctiveness for the COO of HCCs which might be linked to the differential tumor pathology. Overall, our results suggest the predominant COO for the HCCs and Mixed would most likely to be hepatocytes. Also, our evidences point to the possibility of cholangiocytes as a predominant COO for BTCAs, whereas the COOs of ICCs would vary by individual samples. These results implicate the importance of anatomical locations on the possible COOs of PLCs and BTCAs.

**Fig. 2.**
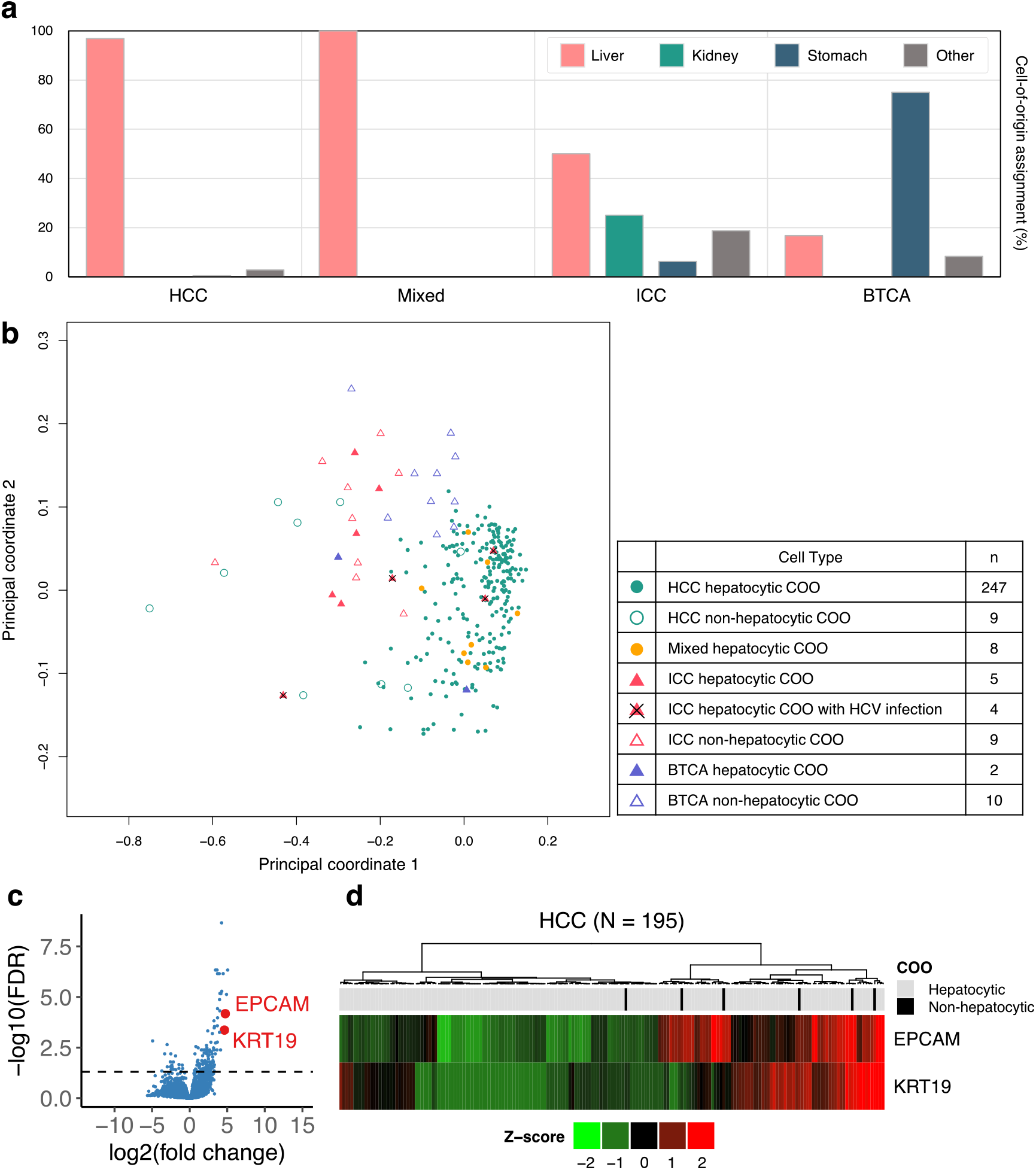
Analysis of COOs at the individual cancer samples. **a** Prediction of COO via grouping of chromatin features for each normal tissue type. Bar graph represents the percentage of samples with respect to the assigned COO by liver tissue chromatin features (pink), kidney tissue chromatin features (green), stomach tissue chromatin features (navy) or the rest (gray). **b** Principal coordinate analysis (PCOA) of mutation frequency distributions for individual cancer samples. **c, d** Differential gene expression by non-hepatocytic COO HCCs (n = 6) comparing to the hepatocytic COO HCCs (n = 189). **c** Volcano plot. The horizontal axis is the log-ratio of the non-hepatocytic COO to the hepatocytic origins. Dashed line represents FDR = 0.05. **d** Expression profile of *EPCAM* and *KRT19* mRNA.

Alongside with these result, principle coordinate analysis (PCOA) result revealed that the PLCs with hepatocytic predicted COO tend to aggregate as a cluster with all of the samples displaying principle coordinate 1 value over 0 (Additional file 1: Figure S4). In terms of the PLC subtypes, HCCs and Mixed samples were all contained within a cluster except for the ones with non-hepatocytic predicted COOs, whereas the ICCs and BTCAs were more spread out (Fig. 2b), reflecting the distinct mutation landscape patterns.

To demonstrate whether HCCs with non-hepatocytic predicted COO have a unique phenotype compared to the hepatocyte-origin HCCs, we analyzed the gene expression profiles. Among the non-hepatocytic- and hepatocyte-origin predicted HCC samples, tumor RNA-Seq data were available for 6 and 189 samples, respectively (4). A comparison of gene expression levels between them showed that 124 genes were up-regulated and 21 genes were down-regulated in non-liver-origin HCCs (FDR < 0.05, absolute logFC > 0.647; Additional file 1: Table S1). Interestingly, the upregulated genes included an epithelial cell marker *EPCAM* and a cholangiocyte-specific marker *KRT19* (Fig. 2c). Clustering analysis confirmed that HCCs with non-hepatocytic predicted COO were enriched in a cluster that expressed more *EPCAM* and *KRT19* (Fig. 2d). Gene set enrichment analysis showed that molecular pathways associated with bile acid synthesis, xenobiotic degradation, and hepatocyte nuclear factor were down-regulated in HCCs with non-hepatocytic predicted COO (Additional file 1: Figure S5). This result indicates that the functional similarity to hepatocyte was lower in HCCs with non-hepatocytic predicted COO compared to hepatocyte-origin HCCs. Collectively, mRNA expression in non-hepatocyte-origin predicted HCCs partly resembled that of biliary epithelial cells. We also compared hepatocyte- and non-hepatocyte-origin predicted HCCs in terms of clinical features (including tumor stage and survival), but we did not see statistically significant difference in these features, implying that the COO assignments for HCCs might be independent from the clinical prognosis.

Previous publication described the association between hepatitis virus infection status and the liver COO assignments without any subgrouping of the virus types (12). As of further investigation, we tested whether there are any hepatitis virus-type dependent tendencies to particular COOs and the associated variance explained scores for the somatic mutation landscape of PLCs. Upon grouping the PLCs with the hepatitis B virus (HBV) and hepatitis C virus (HCV) infection status, our analysis revealed that HCCs and Mixed samples were mostly assigned to hepatocytic predicted COO regardless of the either hepatitis virus infection status. In contrast, COO predictions on HCV-infected ICCs displayed predominance towards hepatocytic predicted COO (n=5, binomial probability of 0.08, two tailed) and HBV infected ICCs mostly displayed non-hepatocytic predicted COO assignments (n=9, binomial probability of 0.04, two tailed) (Additional file 1: Figure S6a, c). Furthermore, spearman correlation values between the regional mutation frequency of aggregated samples grouped by HBV or HCV infection status and the normal liver tissue H3K4me1 chromatin mark level was higher for the HCV-infected ICCs comparing to any other ICCs with different virus infection status, and this result was fully replicated when using the H3K4me1 chromatin marks derived from HBV or HCV-infected liver tissues, thus ensuring more relevancy (Additional file 1: Table S2). In line with these results, variance explained scores for the ICCs calculated by using a total of 9 cell or tissue types, we discovered that the chromatin features with the highest level of variance explained scores were derived from different tissues depending on the hepatitis infection status of ICCs (HBV = kidney tissue, HCV = liver tissue, NBNC = stomach tissue) (Additional file 1: Figure S6b). Albeit limited number of virus infected ICC samples, our results implicate a potential skewness of COO depending on the virus infection status, and a separate cohort level study with larger number of samples is strongly warranted. In addition, these results also reflects the previous findings in differential infectivity of HBVs and HCVs for cholangiocytes (22, 23).

### Hepatic Progenitor Cells as a Possible Cell-of-origin for PLCs

HPCs, so called as oval cells, are a progenitor cell type located inside the Canal of Hering with both hepatocytic and cholangiocytic differentiation capacity and suspected as a possible COO for PLCs. To examine the possibility of HPCs as a possible COO for different subtypes of PLCs, we performed the random forest feature selection analysis using somatic mutation frequency data of HPCs (14) at an aggregate sample level along with the epigenome feature counts. Results from this analysis demonstrated that the mutation landscape of HPCs cannot be explained adequately by the chromatin landscape, with variance explained scores for the top rank chromatin feature and the total 423 features were either below 0 or 25% (Fig. 3a). In contrast, mutation frequency data of colon stem cells (14) (counterpart stem cell type) at an aggregate sample level were explained by pre-existing set of chromatin features with variance explained score over 40% for the H3K9me3 rectal mucosa mark and over 60% for the total 423 features. Post-adjustment of mutation load for colon stem cells at the level of HPCs still showed chromatin marks derived from the rectal mucosa tissue as a top rank feature with over 28% variance explained score, implicating that the differential mutation load might not be a contributing factor for the distinct feature selection analysis results. These results infer distinct mutation landscape between the HPCs and other PLCs, and thus points out the possibility that HPCs might not be a direct COO of PLCs.

**Fig. 3.**
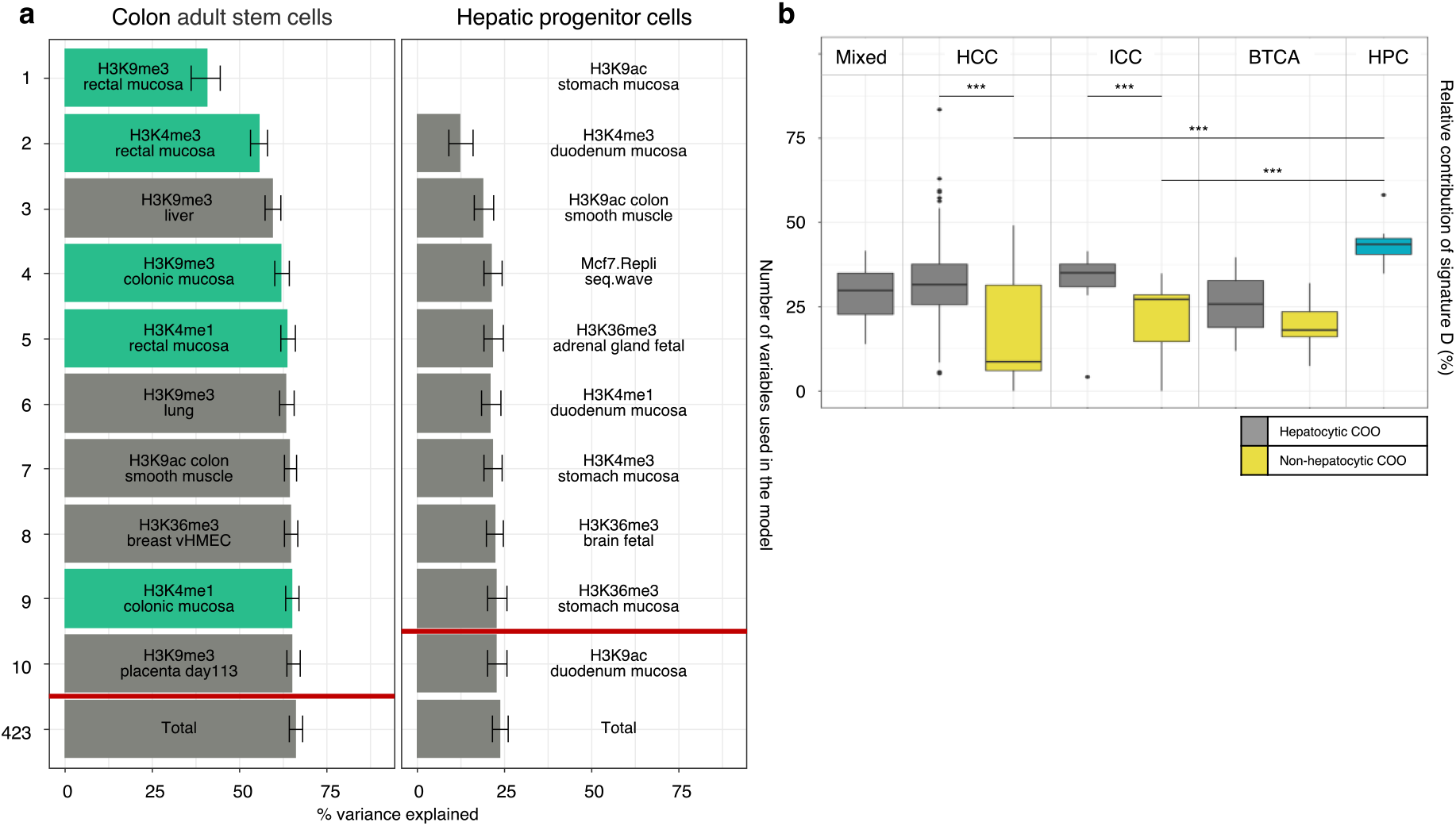
Hepatic progenitor cells have distinct mutation landscape and mutational signature processes compared to primary liver cancer genomes. **a** Chromatin feature selection in relation to the regional mutation frequency of colon adult stem cells and hepatic progenitor cells. Chromatin features related to each tissue type are green-colored. **b** Box plot represents the distribution of relative contribution of signature D in HCC, Mixed, ICC, BTCA and HPC samples. Samples of each tumor type are separated based on whether they are predicted as hepatocytic COO (gray) or not (yellow). Statistical significance is calculated by using Mann-Whitney U-test (***, P < 0.05). BTCAs were excluded from the statistical analysis because only two samples were predicted as hepatocytic COO.

Mutation signature analysis on the somatic mutation landscape of HPCs were previously performed, and identified a specific age-associated mutation signature displaying correlation with the replication timing and the average chromatin level of cell lines registered in the ENCODE project (14). Based on these findings, we conducted the mutation signature analysis on the HPCs along with the PLCs and BTCAs. As predicted, we successfully extracted a resembling signature (signature D) to the age-associated signature previously identified in the HPCs with similar relative proportion level, along with the other three mutation signatures (Additional file 1: Figure S7a, b). Next, we assessed whether the proportion of signature D is correlating with the COO assignment for PLCs. As demonstrated in Fig. 3b, relative contribution level of signature D was significantly lower for non-hepatocytic predicted COO assigned HCCs and ICCs comparing to the hepatocytic predicted COO assigned HCCs / ICCs and all of the HPCs. In line with this, several evidences point out that the correlation between relative proportion of the mutation signature and the COO assignment was specific and consistent for signature D. One is that the proportion of other three signatures (A, B and C) were not significantly associated with the COO assignments for ICCs (P > 0.57) and two signatures (A, B) weren’t showing any any signification associations with the COO assignments for HCCs, too (P > 0.24). Also, mutation type pattern of HPCs were more comparable to the ICCs and BTCAs rather than the HCCs and Mixed, in contrast to the findings on the skewness of COO assignment depending on the signature D status. Furthermore, major proportion of the non-hepatocytic predicted COO samples were located in the lower quartile for the signature D proportions (Additional file 1: Figure S7d). Collectively, these results provide a novel perspective in terms of the possible importance of age-associated mutation signature level on the COO assignment, and thus reflecting again the distinct mutation landscape between the hepatocytic and non-hepatocytic predicted COO samples.

## Discussion

In this paper, we applied random-forest machine learning algorithm and other computational analyses to whole genome sequencing data of PLCs and epigenomics data derived from normal tissues to elucidate unique association patterns between the two features and identify possible cell-of-origin distribution for PLCs at the subtype and individual tumor tissue level. Results from these analyses would help to understand the complex and heterogeneous nature of cancer cell-of-origin and the contribution of chromatin marks on differential regional somatic mutation landscape during the progression for various subtypes of PLCs.

Several recent studies support the idea of chromatin marks serving as a crucial factor in shaping the mutation landscape for several types of tumors (18, 21, 24). Consistent with this idea, our results show that the chromatin marks can explain the mutation landscape of PLCs at the subtype level, displaying variance explained scores in the range of 56% (ICCs) to 87% (HCCs). Moreover, the top chromatin marks associated with the mutational landscape of 256 HCCs were mostly derived from the liver tissue and the top correlative chromatin marks for 12 of BTCAs were from the stomach tissue, which also directly matches to the previous results from the HCCs and DCCs (12). One thing to note is the lower level of variance explained scores for ICCs comparing to any other PLC subtypes. We speculate that the potential contributor to these differences in variance explained scores might be the lower mutation load and the higher level of heterogeneity in COOs at the individual tumor tissue-level since the COOs for individual ICC tissues were the most heterogeneous among all subtypes of PLCs, although following the cell and epithelial types with respect to the anatomical locations of ICCs.

The COOs for PLCs were highly debated for a number of years not only due to the discovery of several types of HPCs (25, 26), but also the facultative regeneration of hepatocytes and cholangiocytes which mainly occurs during the inflammation (7, 11). Our results suggest towards the differentiated cells rather than progenitor or stem cells as origins for PLCs based on the findings that the normal liver (representing hepatocytes), kidney and stomach (surrogate for the cholangiocytes) tissues can mostly explain the COO of PLCs, and the somatic mutation profile from the HPCs is not adequately explained (variance explained score < 24.04) by the normal tissue chromatin marks, albeit the significance of non-hepatocytic predicted COO assignments in regards to the age-associated mutation signature-specific manner. Although our chromatin feature selection analysis did not contain any liver progenitor/stem cell chromatin marks, poor correlation between the mutational landscape of HPCs and the liver or stomach chromatin marks may infer the distinctiveness of chromatin landscape between the differentiated cells/tissues and the progenitor/stem cells. Although we cannot fully reject the possibility that the HPCs are still the very first COO of PLCs, our results at least suggest that the major somatic mutation accumulation would most likely to happen on differentiated cells, not at the progenitor/stem cell level. Future assessment on the relationship between the chromatin marks derived from the HPCs and the mutational landscape of PLCs and HPCs might be a separate confirmatory study, although the limitation on the number of progenitor/stem cells directly from human liver and its purity are major hurdles for ChIP-seq or any other epigenomics assays to be performed.

## Conclusions

In summary, our results on the COO of PLCs discovered several novel and heterogenous nature of COO distributions in different subtypes. We believe that these results address the novel aspects of individual-level differences in tumor biology and clinical pathology of PLCs, and providing a robust and relevant way of studying cancer COO in human system without utilizing a human organoid system, which might be solely suitable for mechanism studies in a practical manner due to the labor intensiveness caused by making each organoid per each patient and potential selection bias during cell culture. Ultimately, our results might add significant arguments for the necessity of personalized medicine for cancer treatments, combined with the genomics and the other molecular signatures.

## Acknowledgements

Not applicable

## Availability of data and materials

WGS and RNA-seq data are deposited in the European Genome-phenome Archive, and these data can be accessed with the approval of the ICGC Data Access Compliance Office. NIH Roadmap epigenomics data can be obtained at Gene Expression Omnibus site (https://www.ncbi.nlm.nih.gov/geo/) under accession number NCBI GSE18927. In addition, chromatin data of liver tissues derived from hepatitis virus infected patients (donor HPC8 and HPC17) can be accessed at IHEC (https://epigenomesportal.ca/ihec/download.html).

## Authors’ contributions

HL provided the original idea. HL, KH, and HK led the overall project. KH, MF, RK, SY, and HL analyzed the data and contributed to scientific discussions. HL and KH wrote the manuscript, and MF, RK, PP, YH, HN, and HK reviewed the manuscript.

## Ethics approval and consent to participate

Not applicable

## Consent for publication

Not applicable

## Additional file

Supplementary tables and figures for Ha et al. “Somatic mutation landscape reveals differential variability of cell-of-origin for primary liver cancer”.

**Table S1.**
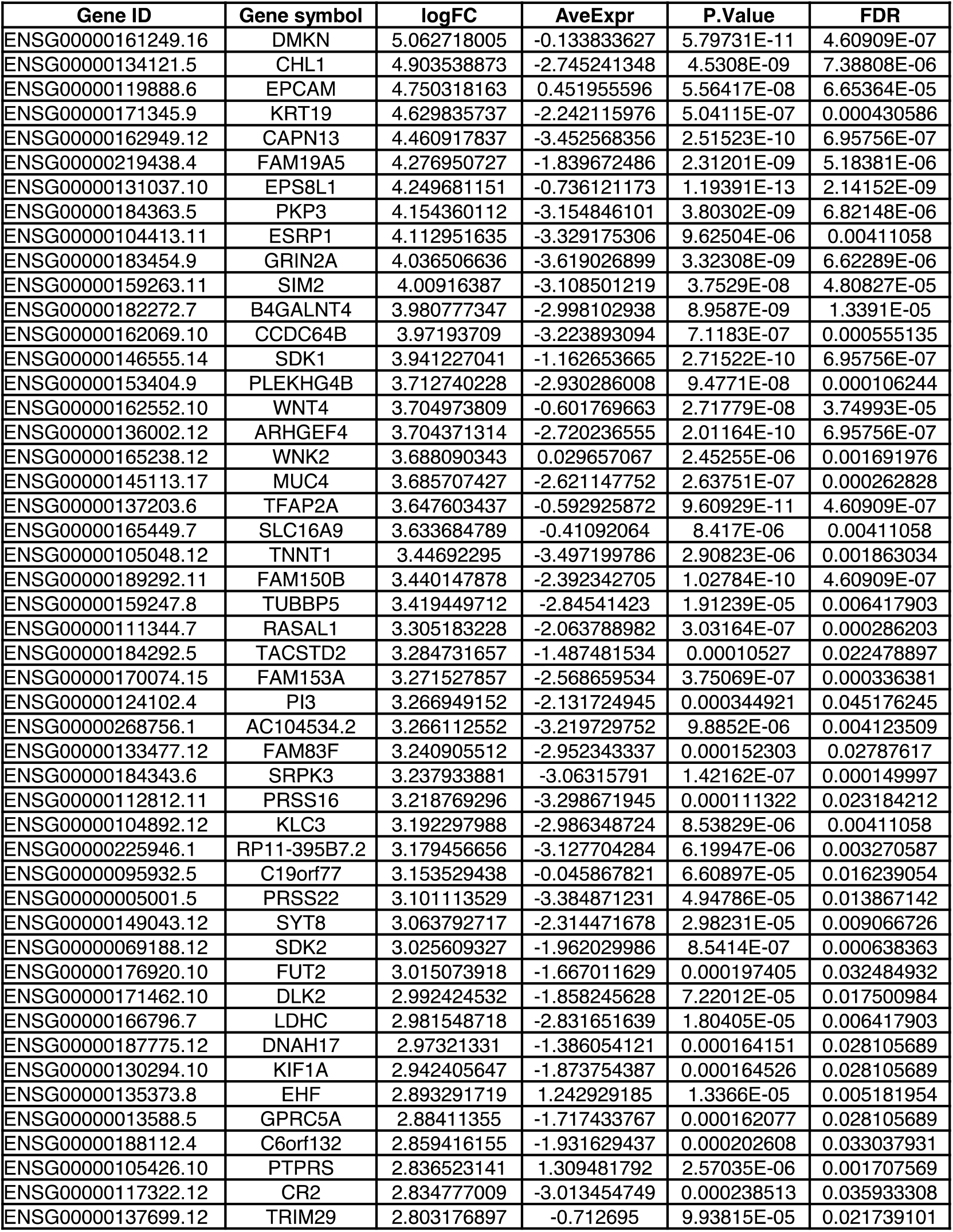
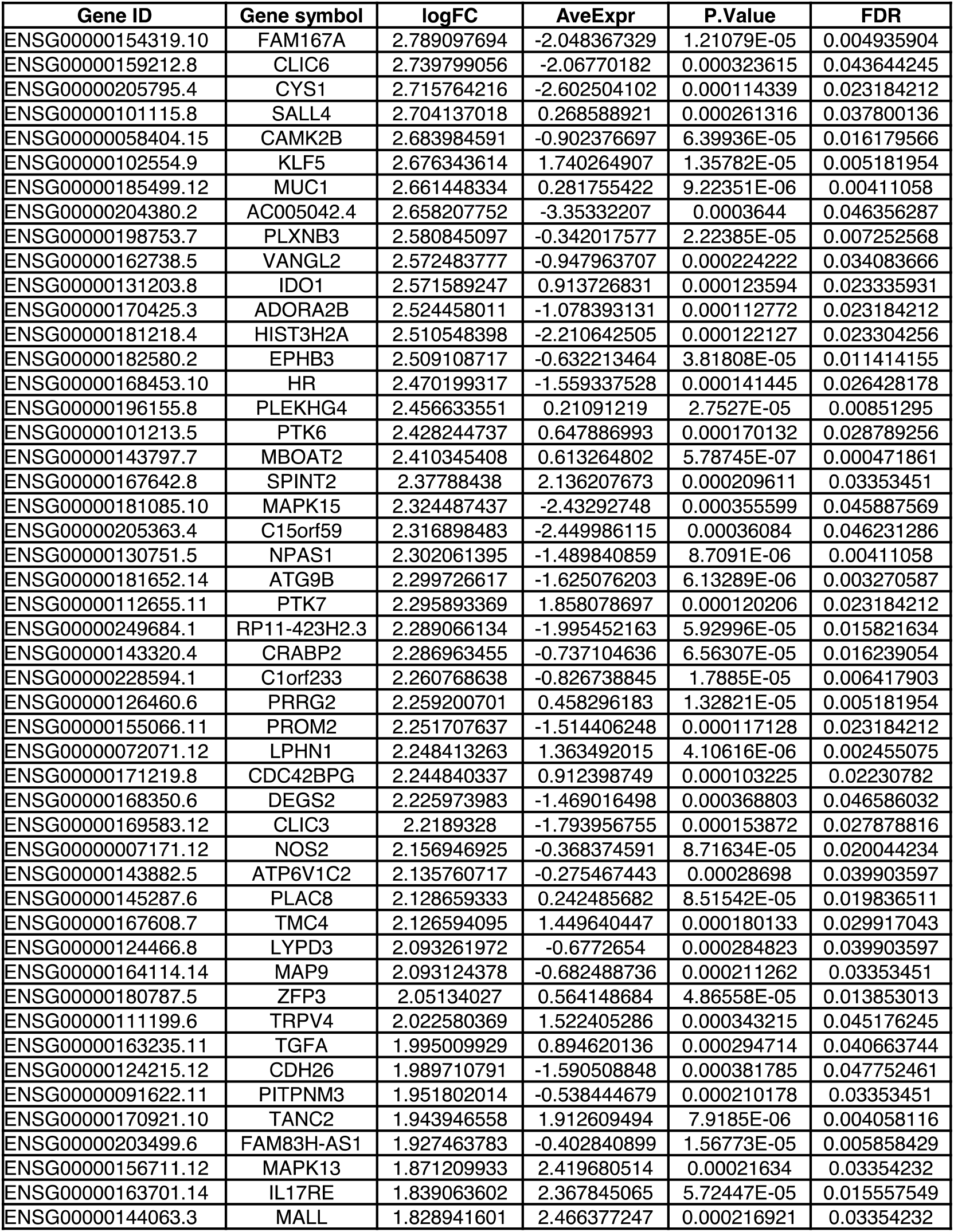
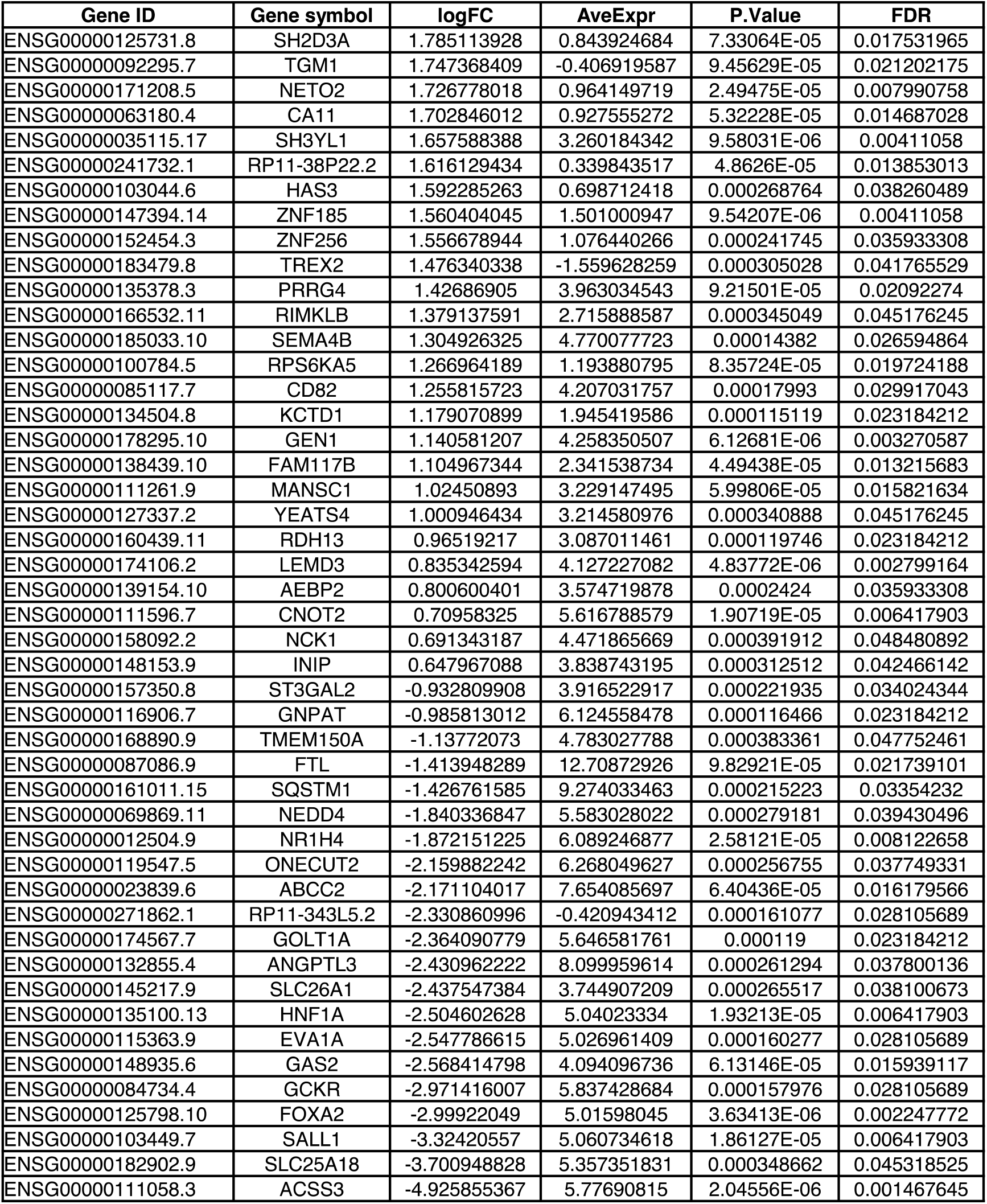
Differentially expressed Genes between non-hepatocytic- and hepatocytic-origin HCCs.

**Table S2.** Spearman correlations between the regional mutation frequency of aggregated sample per infection status of PLCs and level of liver H3K4me1 chromatin mark.

| Spearman's rank correlation abs (r) | HCC |  |  | Mixed |  |  | ICC |  |  |
| --- | --- | --- | --- | --- | --- | --- | --- | --- | --- |
|  | HBV | HCV | NBNC | HBV | HCV | NBNC | HBV | HCV | NBNC |
| H3K4me1 normal liver | 0.869 | 0.890 | 0.875 | 0.628 | 0.744 | 0.562 | 0.361 | 0.609 | 0.504 |
| H3K4me1 HBV liver | 0.798 | 0.811 | 0.800 | 0.587 | 0.679 | 0.514 | 0.334 | 0.540 | 0.452 |
| H3K4me1 HCV liver | 0.743 | 0.752 | 0.744 | 0.550 | 0.635 | 0.479 | 0.297 | 0.493 | 0.416 |

**Figure S1.**
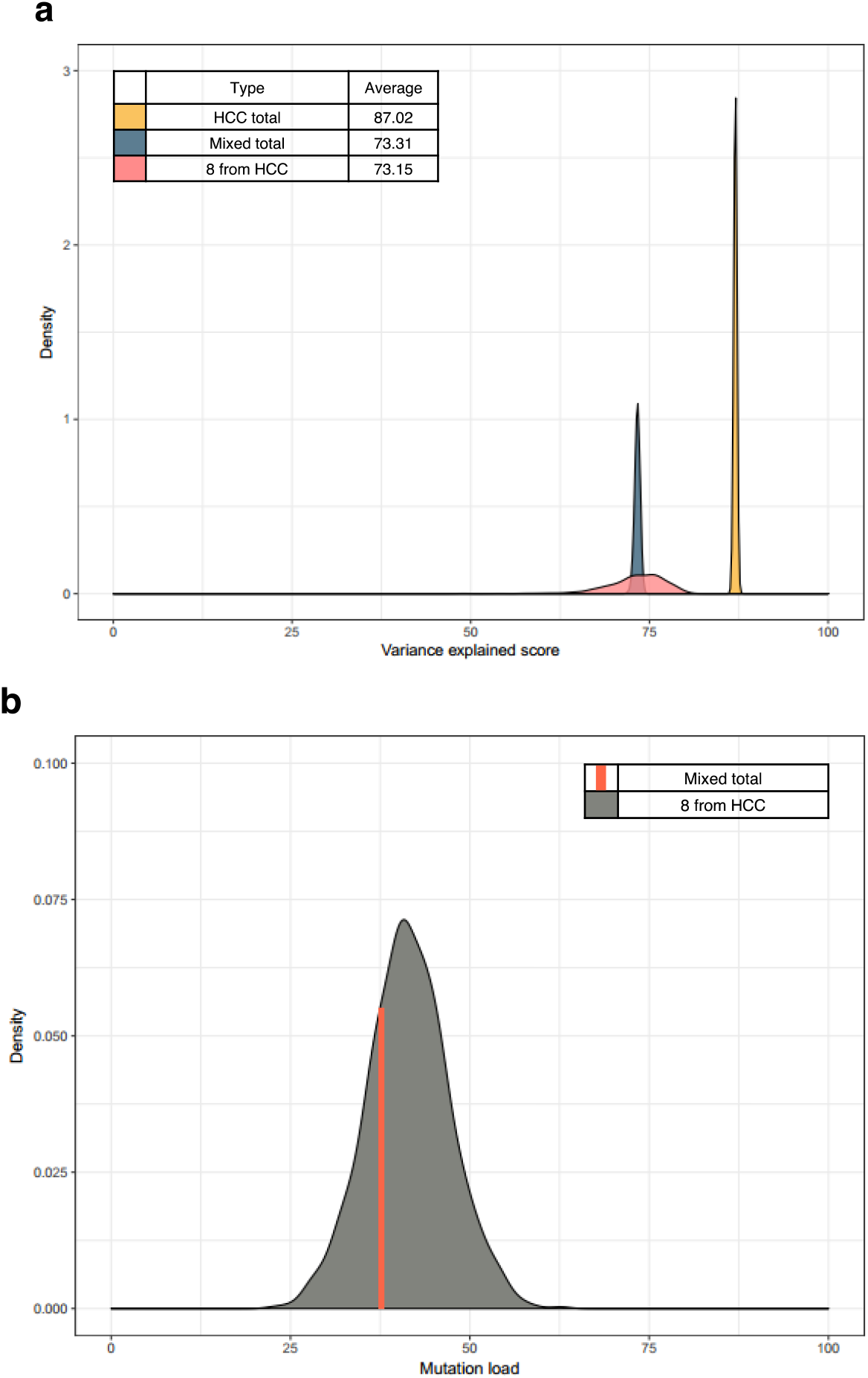
Difference in variance explained scores between the HCC and MIXED type is related to the total number of samples and the aggregated mutation load. (**a**) Distribution of variance explained scores using either all samples or 8 randomly selected samples in 1,000 repeated simulations. Distributions of HCC total (yellow, n = 256) and Mixed total (navy, n = 8) are the result of using all samples for each cancer type. However, pink-colored distribution represents the result of using 8 randomly selected samples in only HCC type. Average variance explained score for each distribution is shown on the top left. (**b**) Distribution of aggregated mutation load at the 1 megabase-level from 8 randomly selected HCC samples in 1,000 repeated simulations. Orange-colored bar represents the aggregated mutation load at the 1 megabase-level from all samples of Mixed type.

**Figure S2.**
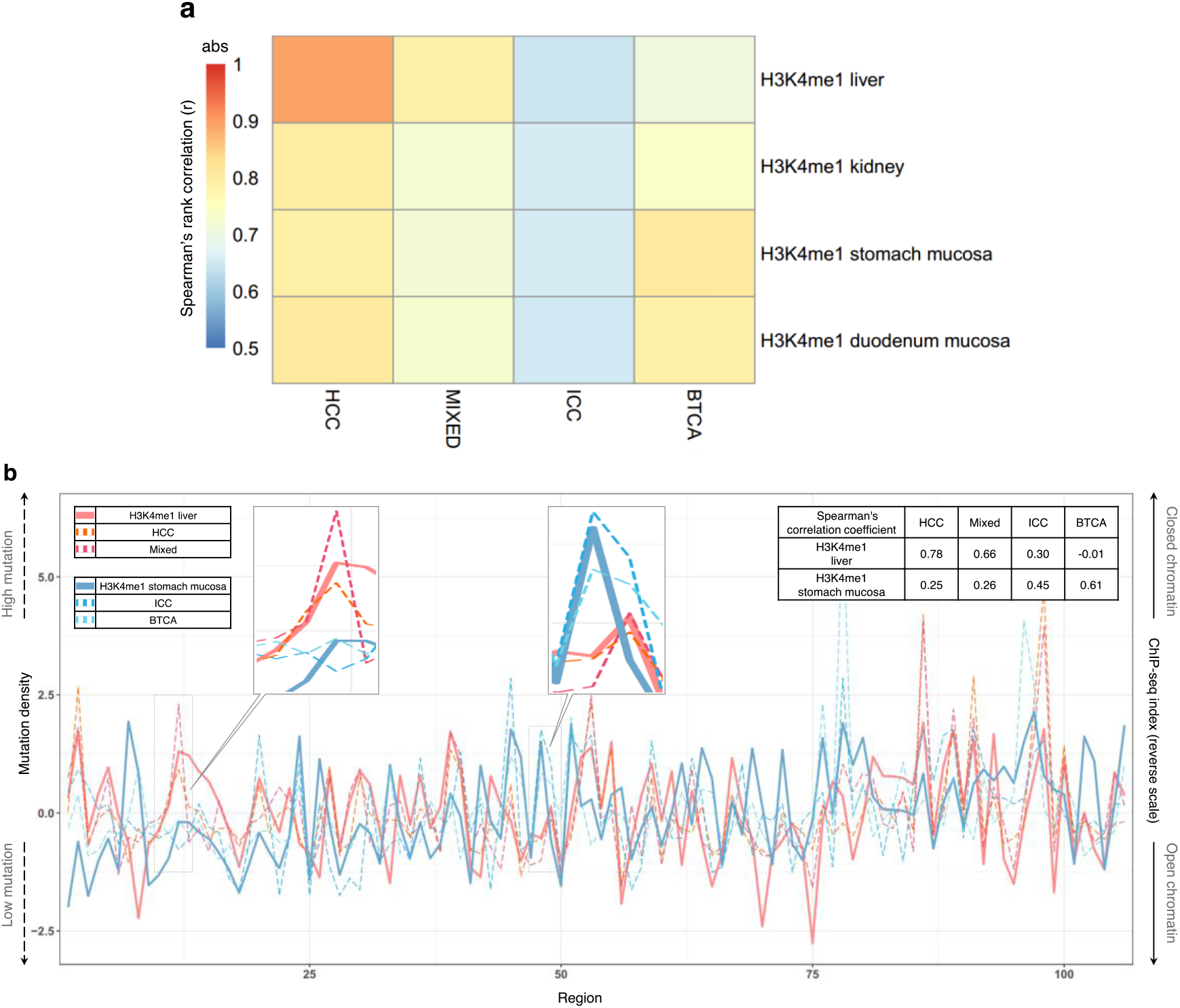
Correlations between cancer genome mutation density and the H3K4me1 chromatin features in different tissue types. (**a**) Heat map with different color depths corresponding to the absolute values of Spearman’s ρ statistics. (**b**) Regional mutation density of HCCs, Mixeds, ICCs and BTCAs parallel to the ChIP-seq index (reverse scale) of liver or stomach H3K4me1. Dotted and solid lines represent mutation density and ChIP-seq index, respectively. A total of 106 genomic regions that show top 5% difference from the predicted ChIP-seq count in the regression model between liver and stomach H3K4me1 were selected. Spearman’s rank correlations between the mutation density and ChIP-seq index are shown on the top right. Zoomed images are representative regions for cancer type groupings with respect to liver and stomach H3K4me1 level (HCC/Mixed and ICC/BTCA).

**Figure S3.**
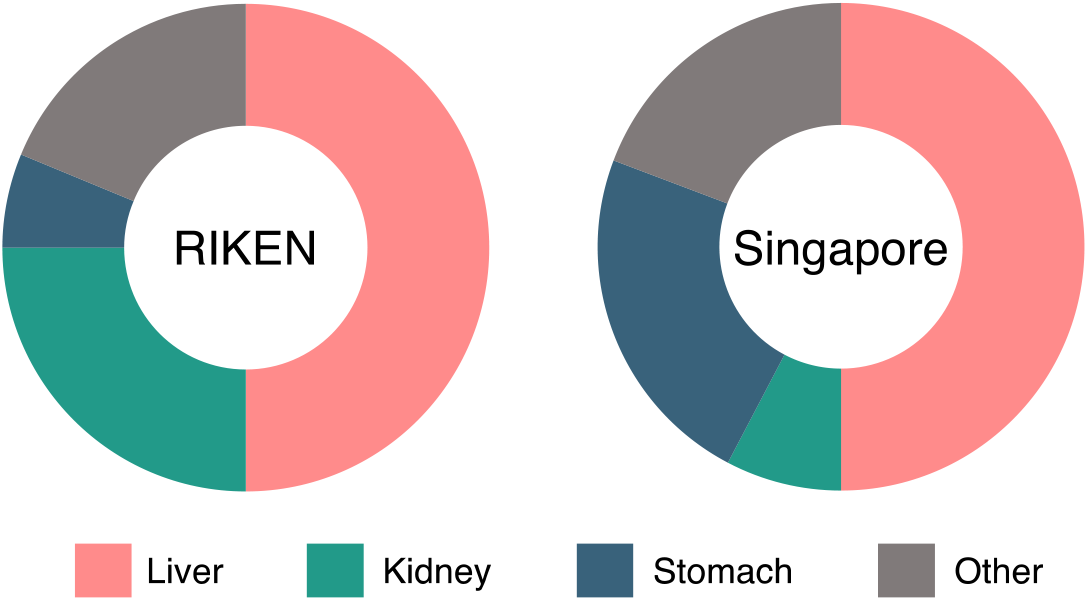
Cell-of-origin prediction distributions for distinct ICC cohorts. Pie graphs represent the percentage of samples getting COO assignments as liver tissue chromatin features (pink), kidney tissue chromatin features (green), stomach tissue chromatin features (navy) or the rest (gray).

**Figure S4.**
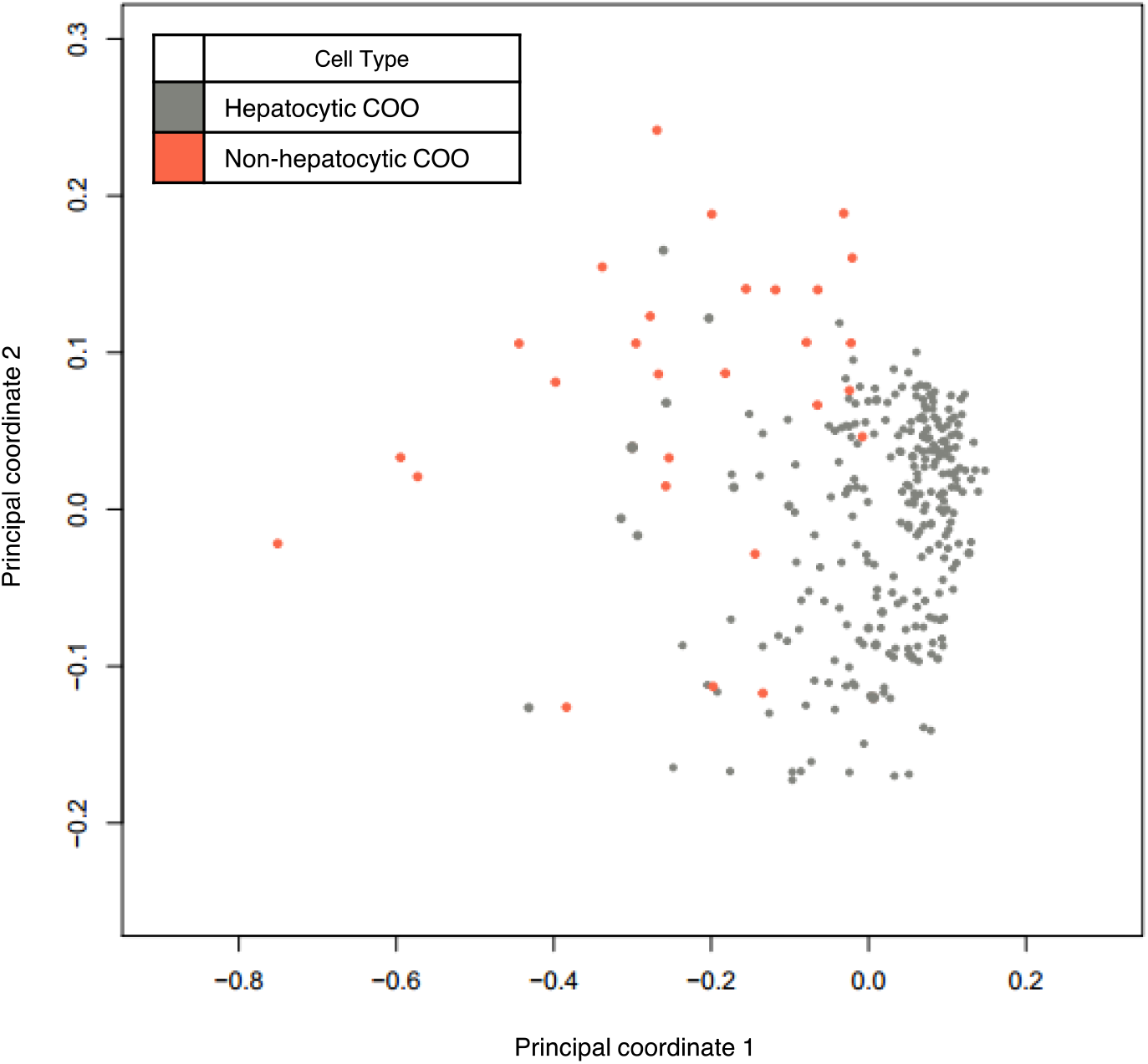
PCOA of individual cancer samples. Hepatocytic COO samples are gray-colored and non-hepatocytic COO samples are orange-colored.

**Figure S5.**
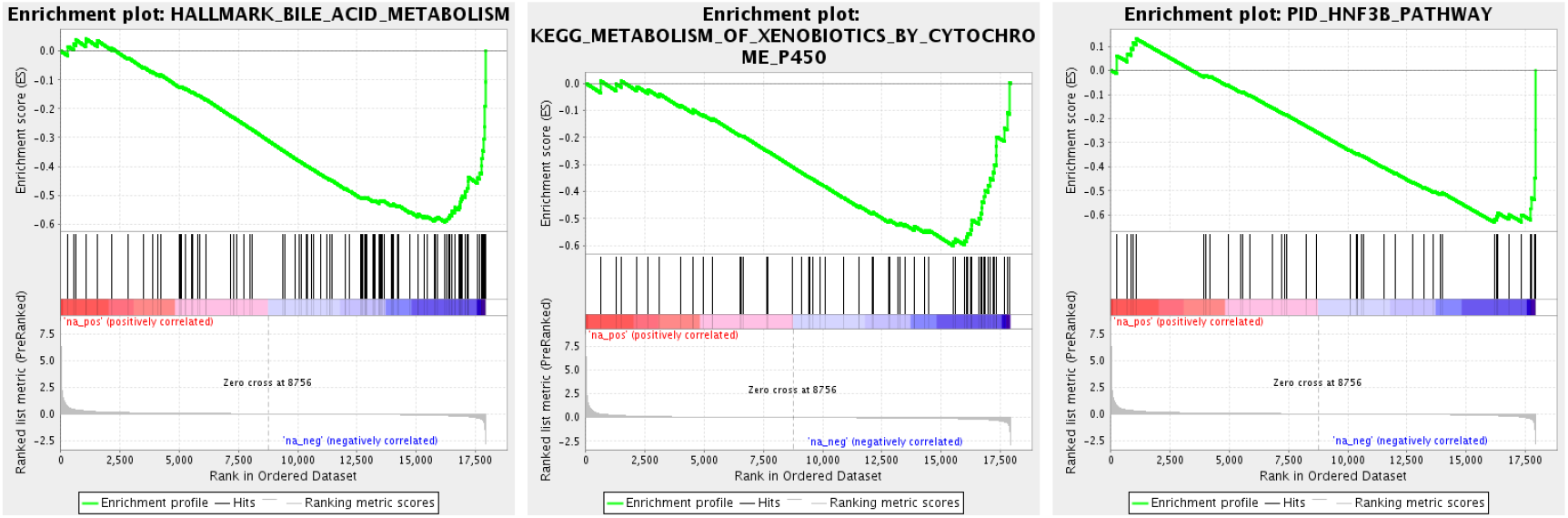
Gene sets that were down-regulated in non-hepatocytic COO HCCs.

**Figure S6.**
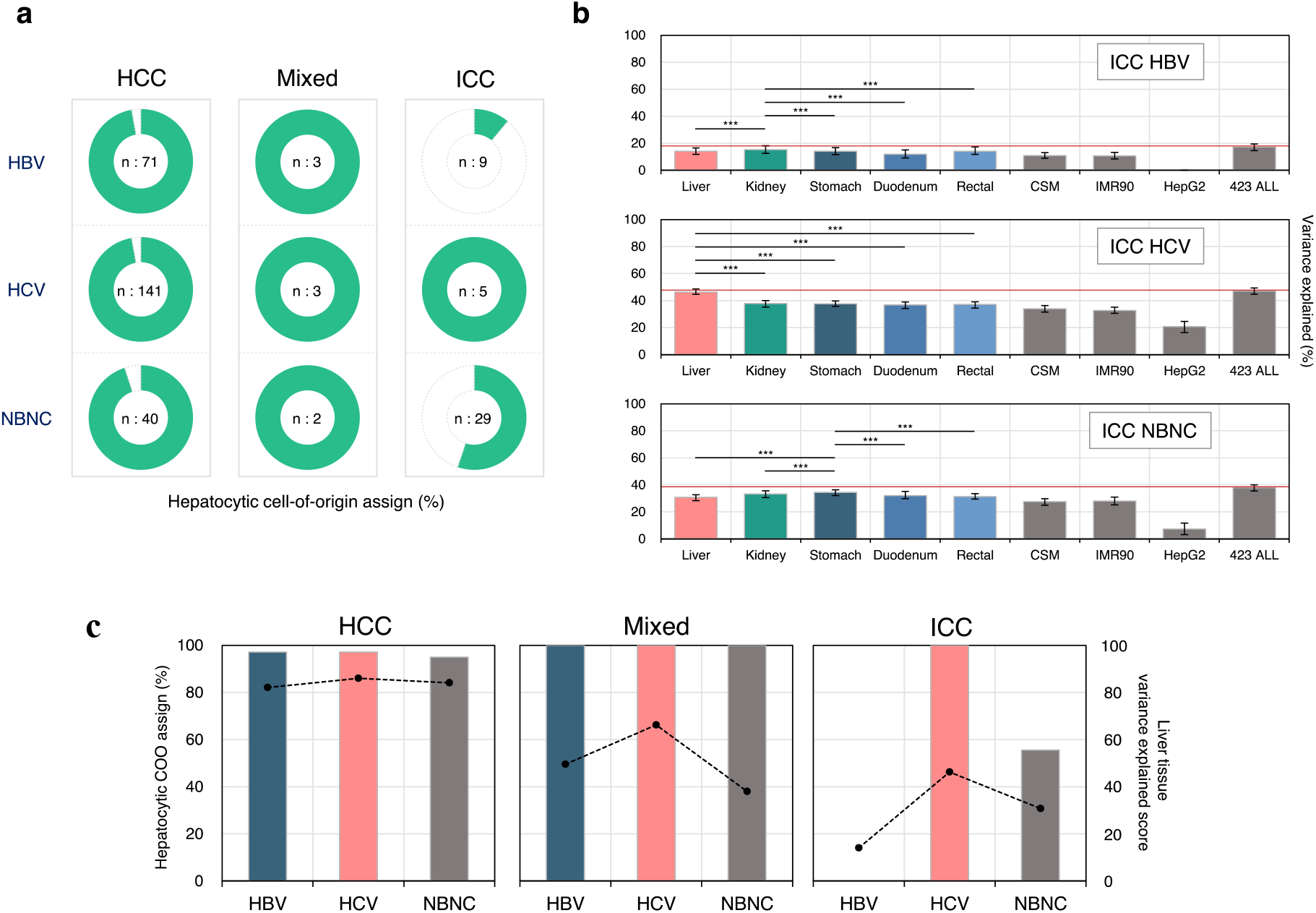
Viral infection status-associated differences in hepatocytic cell-of-origin assignment and variance explained scores. (**a**) Pie graphs represent the percentage of samples assigned as hepatocytic COO (green). The number of samples for analysis is shown at the center of each graph. (**b**) Average variance explained scores for aggregated mutation data depending on the virus type of ICCs are estimated by grouping chromatin features based on each normal cell/tissue type. error bars indicate minimum and maximum scores derived from 1,000 repeated simulations. Red line displays average variance explained score from all 423 epigenomic features. Statistical significance is calculated from tissue with the highest value by using Mann-Whitney U-test (***, P < 0.001). (**c**) Bar graphs show the percentage of samples assigned as hepatocytic COO for RIKEN samples. Dots with lines represent average variance explained scores derived by the liver chromatin features.

**Figure S7.**
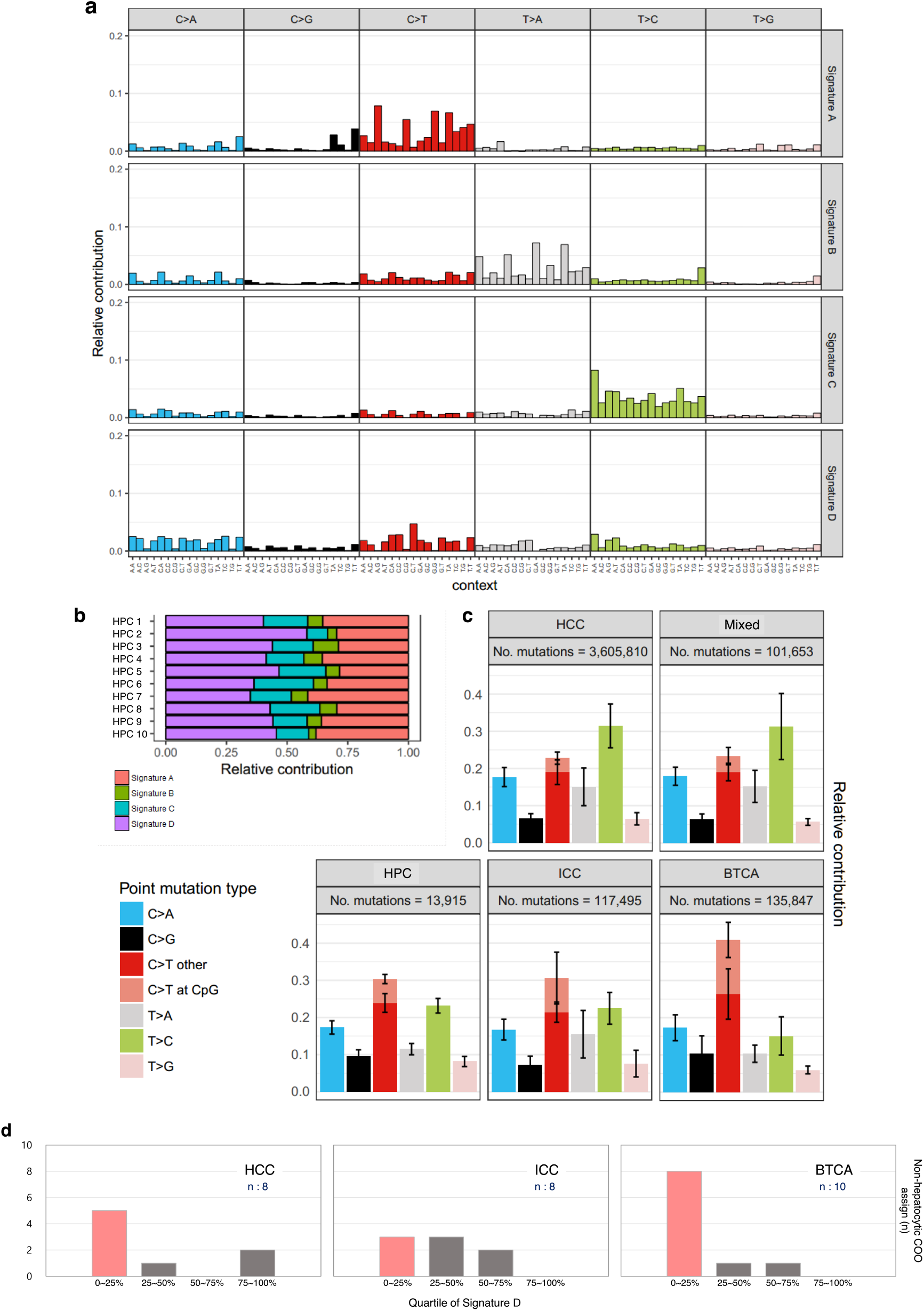
Mutation signature analysis for the genomes of HCC, Mixed, ICC, BTCA-SG and HPC samples. (**a**) Contribution of mutation types to the four mutational signatures derived from the somatic mutations of HCC, Mixed, ICC, BTCA-SG and HPC samples. (**b**) Relative contribution of mutational signatures in each HPC sample. (**c**) Relative contribution of somatic mutation types in each cancer/tissue type. Bar length is calculated as the average relative contribution in each type and error bars show standard deviation. (**d**) Cell-of-origin assignment status based on mutational signatures for HCC, ICC and BTCA. The bar represents the number of non-hepatocytic COO assigned samples with respect to the quartile of signature D contribution. Quartile values are determined by sorting samples of HCCs, Mixed, ICCs, BTCAs and HPCs according to the relative contribution of signature D. The number of samples used in the analysis is shown on each plot.

## Abbreviations

PLC: : primary liver cancer
COO: : cell-of-origin
HCC: : hepatocellular carcinoma
Mixed: : mixed hepatocellular and cholangiocarcinoma
ICC: : intrahepatic cholangiocarcinoma
HPC: : hepatic progenitor cell
BTCA: : extrahepatic biliary tract cholangiocarcinoma
WGS: : whole-genome sequencing
LINC-JP: : NCC-Japan liver cancer
LIRI-JP: : RIKEN-Japan liver cancer
BTCA-SG: : Singapore biliary tract cancer
NMF: : nonnegative matrix factorization
GSEA: : gene set enrichment analysis
DCC: : extrahepatic distal cholangiocarcinoma
PCOA: : principle coordinate analysis
HBV: : hepatitis B virus
HCV: : hepatitis C virus

